# Sex differences in contextual fear expression are associated with altered medial prefrontal cortex activity

**DOI:** 10.1101/2024.09.07.611834

**Authors:** Katherine Vazquez, Ryan G. Parsons

**Affiliations:** Stony Brook University, Department of Psychology, 100 Nicolls Rd., Stony Brook, NY, 11794

**Keywords:** contextual Fear, sex difference, infralimbic, prelimbic, periaqueductal gray

## Abstract

Understanding the neural basis of fear expression in rodents has implications for understanding pathological fear responses that characterize posttraumatic stress disorder. Even though posttraumatic stress disorder is more common in females, little is known about the neural circuit interactions supporting fear expression in female rodents. In this study, we were interested in determining whether neural activity associated with the expression of contextual fear differed between males and females within the projections from the medial prefrontal cortex to the ventrolateral periaqueductal gray, and in the medial prefrontal cortex in neurons that do not project to the periaqueductal gray. We infused a viral retrograde tracer into the ventrolateral periaqueductal gray in male and female rats and trained them in a contextual fear conditioning task. The following day rats were re-exposed to the conditioning context and were sacrificed shortly thereafter. Neural activity was measured using EGR1 immunofluorescence. The behavioral results showed that males exhibited higher levels of freezing during the context test than females. Male rats that underwent training and testing showed an increase in the proportion of viral infected cells that express EGR1 in the PL compared to rats that had only received context exposure. Trained female rats were not different than controls, however a direct comparison between sexes was not different. In cells not labeled by the tracer, males showed higher levels of fear-induced EGR1 expression in the prelimbic cortex than females. Conversely, females showed higher levels of EGR1 expression in the infralimbic cortex following testing as compared to males. These results suggest that sex differences in the expression of contextual fear may involve differences in the relative activity levels of the prelimbic and infralimbic cortex.

## Introduction

Included in the core features of post-traumatic stress disorder (PTSD) are excessive fear to trauma-associated cues and the generalization of fear behavior to stimuli not associated with a traumatic experience. Therefore, determining how organisms detect and respond to threats, and how this process becomes dysregulated, is of vital importance. Much of the prior work looking into the behavioral and neural mechanisms of fear reactions has used Pavlovian fear conditioning, a fundamental form of learning where organisms acquire relationships between aversive stimuli and the cues that predict them. Through the study of Pavlovian fear conditioning in rodents, many components of the neural systems regulating the expression of fear have been identified [1,2]. However, even though females show much higher rates of PTSD, most of the basic research on fear expression in rodents has been completed using only male subjects. Given this, it is essential that studies of fear learning and expression in rodents use both sexes.

Many studies in rodents have reported that females show reduced contextual fear compared to males [3–7], although this effect is dependent on several factors [8–11]. In some studies that have observed sex differences in contextual fear, the differences have been shown to be associated with sex differences in hippocampal plasticity [3, 4, 12], possibly reflecting sex differences in spatial learning.

There is also evidence that regions within the medial prefrontal cortex (mPFC) play a key role in contextual fear conditioning and the expression of contextual fear. These include the prelimbic cortex (PL) which previous studies have outlined a role for in the acquisition and expression of both cued and contextual fear [13–15], and the infralimbic cortex (IL) which has most frequently been associated with the extinction of fear learning but has also been implicated in the acquisition and expression of contextual fear [16, 17]. Although some studies have described sex differences in the function of IL and PL between males and females as it relates to fear processing [18–21], whether sex differences in contextual fear involves alterations in mPFC function is not known.

Several studies have described how interactions centered on the mPFC contribute to various processes related to fear learning and expression [22–28]. However, little is known about the function of projections from the mPFC to the periaqueductal gray (PAG), even though both brain regions are known to be involved in the expression of fear behavior [13, 29, 30]. One study [31] showed that projections originating in the dorsal region of mPFC (i.e. prelimbic cortex and anterior cingulate cortex) to the ventral and lateral regions of the PAG are involved in the discrimination, but not expression, of contextual fear. However, this study used only males as subjects and was not able to distinguish the contribution of specific regions within the PAG or PFC.

The goal of the present study was to assess neural activity related to the expression of contextual fear in the mPFC projections to the vlPAG in both male and female rats. Additionally, we also wanted to measure neural activity more broadly in the mPFC given how few studies have compared neural activity following fear expression in males and females. We infused a viral retrograde tracer into the vlPAG of rats of both sexes and exposed them to contextual fear conditioning and testing. We assessed neural activity following the test session by counting the number of EGR1 positive cells in the anterior cingulate, prelimbic, and infralimbic cortices both in the cells labeled by the retrograde tracer, as well as from cells not labeled by the tracer.

## Materials and Methods

All procedures were conducted with approval from the Stony Brook University Institutional Animal Care and Use Committee and in accordance with the National Institutes of Health guidelines for the care and use of laboratory animals.

### Subjects

Sprague Dawley rats (13 males and 14 females) obtained from Charles River laboratories served as subjects. Rats were housed in pairs in a colony room maintained on a 12 hr light/dark cycle, and food and water were provided freely throughout the experiment. Upon delivery, rats were left undisturbed for 7 days, and then each rat was gently handled for 5 minutes every day for the 3 days prior to surgery. After recovery, rats were handled for 6 days for 5 minutes each. During the last 3 days of handling, rats were carted into the laboratory to acclimate them to being transported. Behavioral procedures began after the sixth day of handling.

### Surgical Procedures

Rats were anaesthetized with either ketamine (87mg/kg) and xylazine (10mg/kg) or Isoflurane (5.0%) and given unilateral infusions of a retrograde transducing adeno-associated virus (AAVretro-CAG-GFP Addgene #37825; [32]) into the vlPAG (for males: AP =-7.6, ML =+/-0.8, DV= -6.2, for females: AP = - 7.6, ML =+/-0.8, DV = -5.7). The hemisphere was counterbalanced across groups. To deliver the virus, a 22-gauge cannula was lowered into place and an internal cannulae (28 gauge) was inserted through the guide cannula. The internal cannulae were connected to PE-20 tubing, which was connected to an infusion pump. The virus (0.4 μl/site) was injected at a rate of 0.15 μl/min, and the internal cannulae remained in place for 5 minutes after the infusion. Rats were given subcutaneous injections of Meloxicam (1mg/kg) after the procedure to relieve pain, and glycopyrrolate (0.02mg/kg, SC) to prevent congestion.

### Apparatus and contextual fear conditioning

Training and testing for contextual fear conditioning were conducted in 32 cm x 26 cm x 21 cm conditioning chambers and freezing behavior was scored by FreezeScan 2.00 Software (Clever Sys) using motion parameters that closely matched hand scored data by a trained observer. Additional details of the apparatus are outlined in [33]. One group of rats were placed into the conditioning chambers and after 6.5 minutes were given 4 unsignaled shocks spaced 30 seconds apart, and the rats were removed after being in the chamber for 10 minutes. A second group of rats were placed in the same chambers for 10 minutes but were not exposed to shock. Twenty-four hours later, all the animals were placed in the same chambers for 10 minutes to measure the expression of contextual fear.

### Histology

Sixty minutes after the start of the testing session, all rats were anesthetized with an IP injection of fatal plus solution (100 mg/kg) and transcardially perfused with ice cold PBS followed by buffered formalin. The brains were removed and stored in a 30% sucrose-formalin solution for at least 48 hours, and then sectioned at 40μm using a cryostat. One set of sections from the vlPAG was taken to assess retrograde labeling, and mPFC sections were processed for immunofluorescence.

### Immunofluorescence

Sections from all animals were processed for immunofluorescence for the early growth response protein 1 (EGR1), an immediate early gene and marker of neural activity. Details of the procedure can be seen in [33]. Briefly, free-floating sections were washed, blocked, and incubated overnight at 4C in primary antibody for EGR1 (Cell Signaling #4153). The next day slices were left at room temp for 30 minutes, washed, and exposed to a secondary antibody (Alexa Fluor-594, anti-rabbit, Life Technologies) for 2 hours at room temperature. Slices were washed, mounted on slides, allowed to dry at room temperature for approximately 15 minutes and then cover slipped using Fluoromount G (Southern Biotechnology). Images were taken from the hemisphere ipsilateral to the injection site using an Infinity 3 camera (Lumenera Corporation) mounted to a Zeiss Axioskop 2 at 10x magnification. Images were digitized using Infinity Analyze software (v6.5, Lumenera Corporation), with identical gain and exposure time settings for all acquired images.

### Data Analysis

#### Behavior

For training, the percentage time engaged in freezing behavior was averaged for each minute of the 10-minute session. These data were then subjected to a mixed factors ANOVA, with minute as a within subject factor, and condition (i.e. trained vs context controls) and sex as the between subject factors. For testing, freezing behavior over the course of the entire 10-minute session was averaged and a Two-way ANOVA with sex and condition as factors was performed.

#### Immunofluorescence

Digitized images were opened in Image J (NIH), and a brightness-contrast adjustment was performed for all images (minimum displayed value – 35, maximum displayed value – 155). A sample box was drawn for each of the regions of interest, and single- and double-labeled neurons were counted for tracer-labeled and EGR1 positive cells, and these counts were corrected for the area sampled (i.e. mm^2^). For the analysis of dual-labeled cells, the proportion of tracer labeled cells positive for EGR1 was computed for each animal. These values were then normalized to the average value of the control animals for each sex and Mann-Whitney U tests were used to compare for differences between condition and sex. Mann-Whitney U tests were used instead of the ANOVA because the cell count data were normalized to controls, making parametric tests inappropriate. For cell counts in neurons not labeled by the tracer, tallies were taken from the same images and adjusted for the size of the sample area. These values were then normalized as described above and analyzed in the same manner. Finally, we also computed the ratio of PL activity normalized to controls to normalized IL activity for each rat in the trained groups and compared males and females using a Mann-Whitney U test.

## Results

Rats with injections of AAVretro-GFP were exposed to a contextual fear conditioning task in which unsignaled presentations of shock were delivered and the expression of contextual fear was tested during a 10-minute session the next day (Figure 1A). Controls were given an equivalent amount of context exposure, but no shock was administered during the conditioning session. We assessed freezing behavior during the fear conditioning (Figure 1B) and testing (Figure 1C) sessions. During training, we used a repeated measures ANOVA with time as a within-subject factor, and sex and condition as between-subject factors. Results from this analysis revealed a significant effect of time (F (9, 230) = 101.1, P<0.0001), a significant effect of condition (F (1, 230) = 348.8, P<0.0001), and a significant time x condition interaction (F (9, 230) = 86.44, P<0.0001). These results indicate that fear levels increased during conditioning in rats that were shocked. During testing, a Two-way ANOVA with sex and condition as factors showed a significant effect of condition (F (1, 23) = 46.31, P<0.0001), revealing that freezing levels during testing were higher in trained rats compared to controls. There was also a main effect of sex (F (1, 23) = 9.726, P<0.01) and a significant interaction of sex and condition (F (1, 23) = 5.588, P<0.05), indicating that females showed reduced expression of contextual fear.

**Figure 1.**
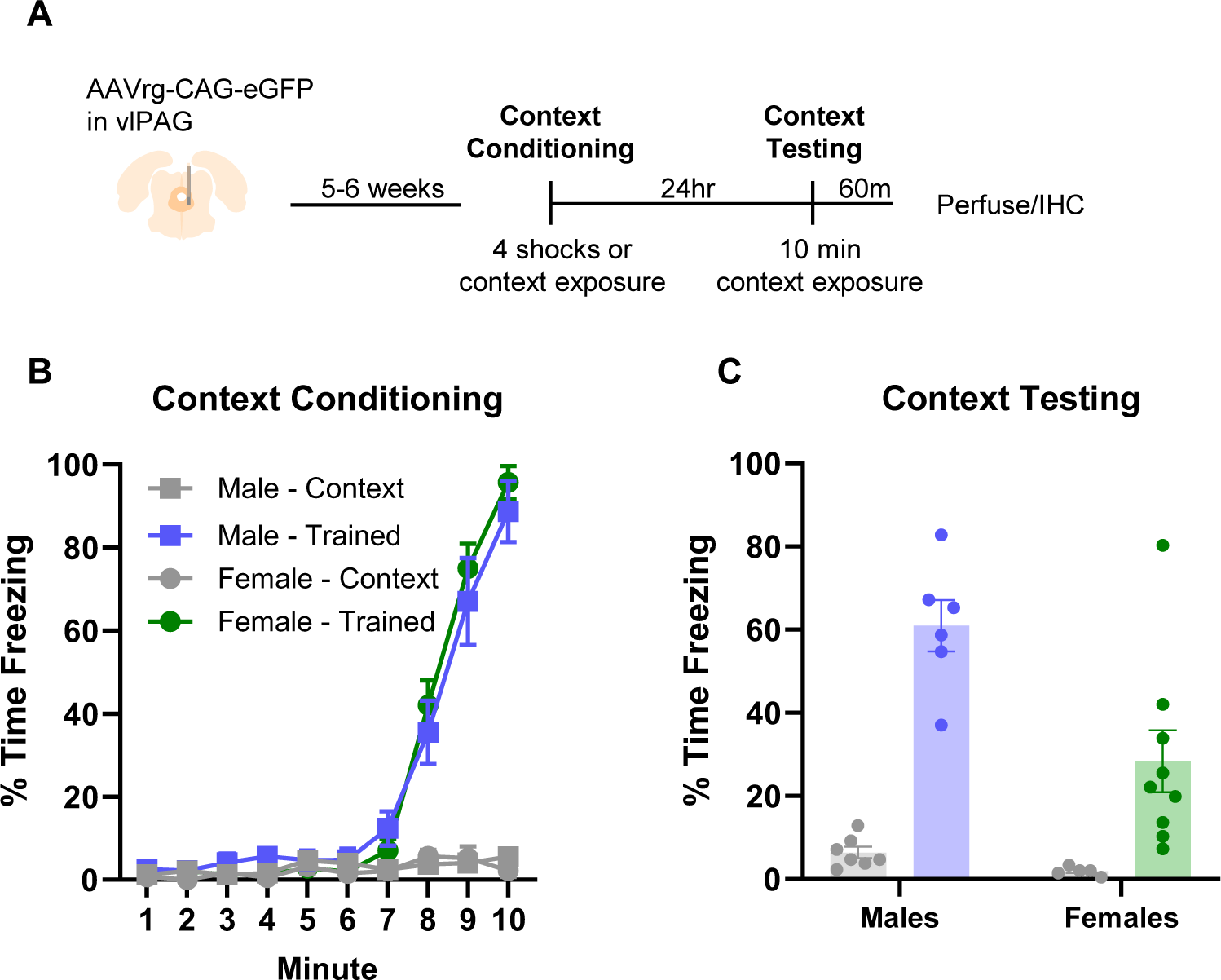
(A) Rats were injected unilaterally with AAVretro-CAG-GFP targeting the vlPAG and after several weeks were given a single contextual conditioning procedure followed the next day by a 10- minute context test. Freezing behavior is depicted during the conditioning session (B) and during the context test the next day (C).

Next, we analyzed the number of GFP-positive neurons in the anterior cingulate (ACC), prelimbic cortex (PL), and infralimbic cortex (IL) in rats that were given AAVretro-GFP into the vlPAG. Figure 2 shows the location and extent of AAV expression in the vlPAG for male (Figure 2A) and female (Figure 2B) rats. Figure 2C depicts sample images indicating the maximum and minimum extent of expression at the injection site. Figure 2D depicts an image showing the distribution and extent of retrograde labeling in the ACC, PL, and IL for a representative subject. Figure 2E shows the average number of GFP-labeled cells in the ACC, PL, and IL for both sexes. We used an ANOVA with sex and subregion as factors to analyze these data. Results from this analysis revealed a significant effect of subregion (F (2, 78) = 9.311, p < .01), but no effect of sex (F (1, 78) = 0.02377, p > 0.05) and no interaction (F (2, 78) = 3.069, p > 0.05). Tukey post hoc comparisons indicate that there was significantly more retrograde labeling in the PL compared to both the ACC (p<0.01) and IL (p<0.05).

**Figure 2.**
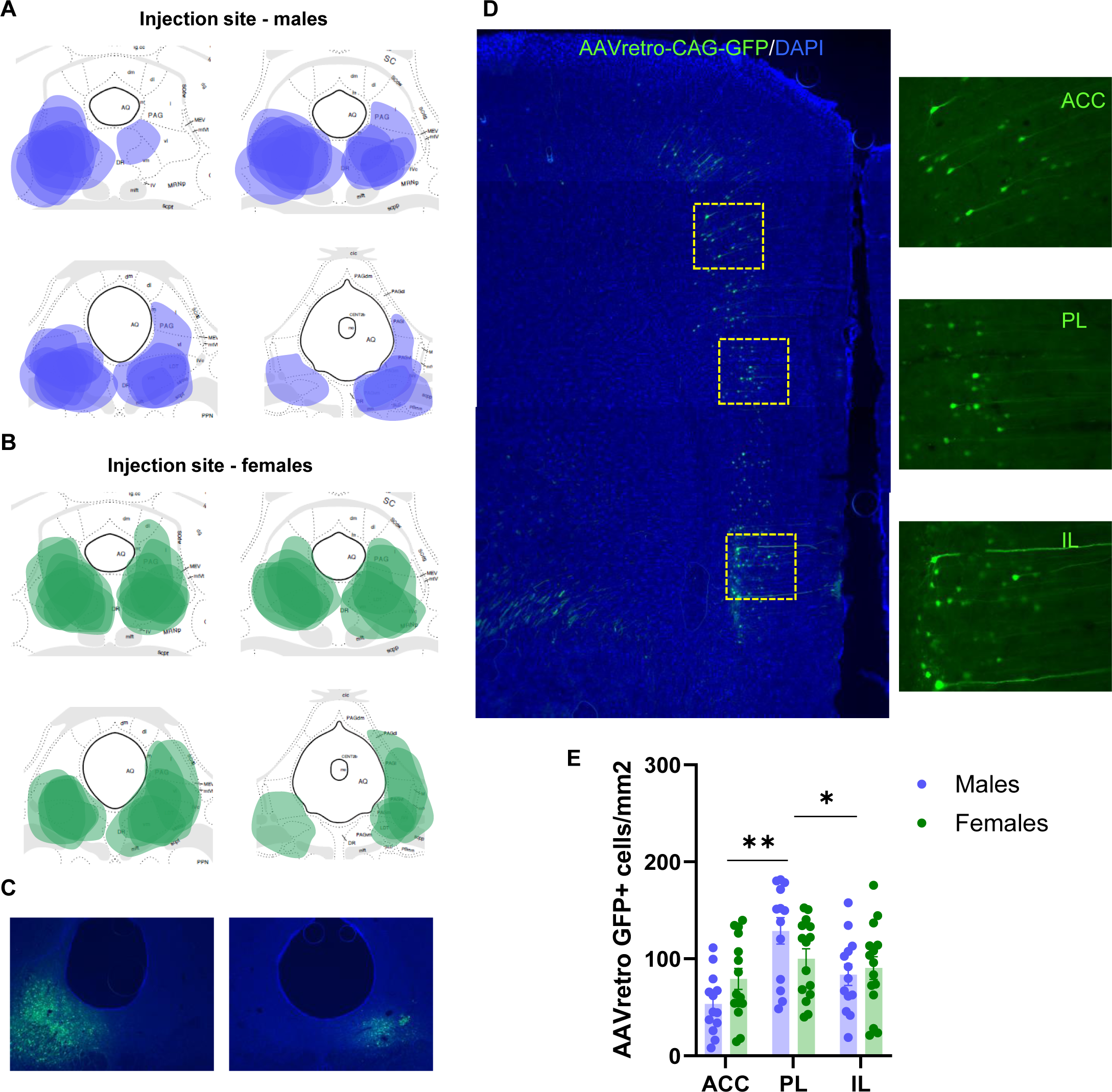
Depiction of injection sites in male (A) and female (B) rats given AAVretro-GFP into the vlPAG. Sample images (C) from two rats showing the largest (top) and smallest (bottom) levels of expression from rats included in the analysis. (D) A representative stitched image (left) at 5x depicting retrograde labeled cells across the medial prefrontal cortex following vlPAG injection of AAVretro. The right side shows higher magnification (20x) images from the ACC, PL, and IL of the same section taken from the yellow outlined boxes on the stitched image. (E) The mean number of retrograde-labeled cells in the ACC, PL, and IL from male and female rats collapsed experimental conditions. ** p < 0.01, * p < 0.05.

In Figure 3 we analyzed the expression of the neural activity marker EGR1 in AAVretro-GFP-labeled cells in the ACC, PL, and IL. Figure 3A displays representative images across all groups showing EGR1 staining (red), AAVretro-GFP (green), and dual labeled cells (orange). Mann-Whitney U tests were used to compare the number of dual labeled cells in trained animals versus within-sex controls and to compare trained male and female animals. When we analyzed EGR1 counts in AAVretro-GFP cells in the ACC (Figure 3B), there was no effect of training in males (U = 20.50, P> 0.05) or females (U = 20.00, P> 0.05), and no difference between males and females (U = 22.00, P> 0.05). Analysis of data in male rats from the PL revealed a significant effect of training (U = 3, P < 0.05; Figure 3C). In female rats there was no significant difference between trained and control rats (U = 14, P > 0.05). In a direct comparison between male and female rats, there was no significant difference (U = 19.00, P> 0.05). Thus, while there was no effect of training in female rats, there was also no difference between sexes. Finally, we analyzed data for EGR1 activity in AAVretro-GFP-labeled cells in the IL (Figure 3D). We found that there was no effect of training in males (U = 19.00, P> 0.05) or females (U = 16.00, P> 0.05), and no difference when trained males and females were compared directly (U = 24.00, P> 0.05).

**Figure 3.**
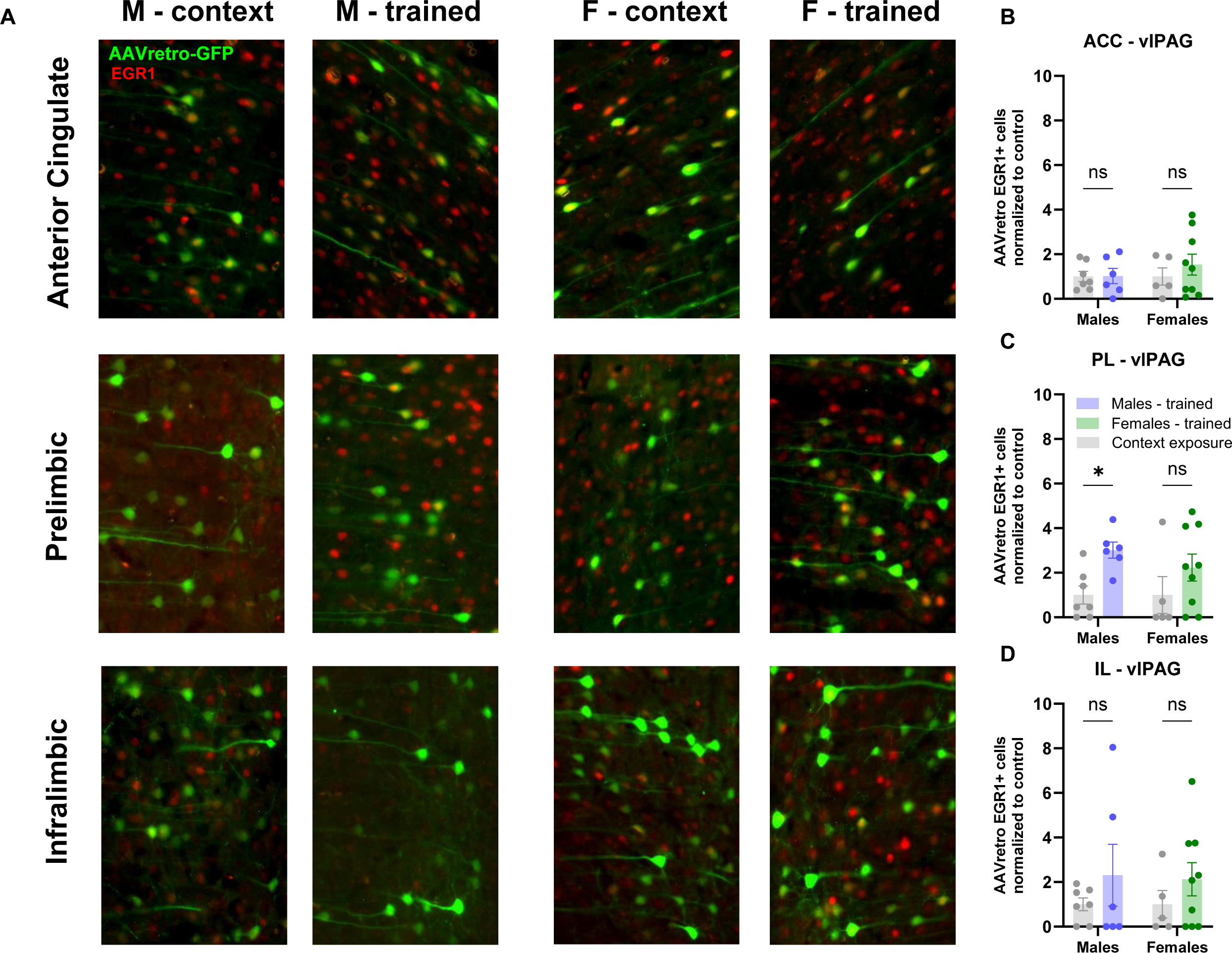
(A) Representative images showing the expression of AAVretro-GFP (green) and EGR1 (red) and dual labeled cells (orange) in the anterior cingulate, prelimbic, and infralimbic regions of the mPFC in males and females. The proportion of dual label red cells in the ACC (B), PL (C), and IL (D) in the different experimental conditions. * p < 0.05

Next, we analyzed the expression of EGR1 in the mPFC in cells that were not labeled with the retrograde tracer (Figure 4). We analyzed the number of EGR1+ cells in trained animals compared to the within-sex controls, and EGR1 expression in trained males versus trained females, using Mann-Whitney U tests. No significant differences were observed in the ACC in males, (U = 21.00, P> 0.05) or females (U = 22.00, P> 0.05), or between males and females (U = 27.00, P> 0.05; Figure 4A). In the PL (Figure 4B) there was a significant difference between trained males and male controls (U = 5.00, P< 0.05), but no difference between females and controls (U = 17.00, P> 0.05). The comparison between male and female rats that were trained was also significant (U = 17.00, P> 0.05), with males showing higher levels of EGR1 expression than females in the PL. Next, we performed the same analyses on data from the IL (Figure 4C). We found that there was no difference between male (U = 14.00, P> 0.05) or female (U = 14.00, P> 0.05) rats and their within-sex controls, however the comparison between males and females that were conditioned was significant, with females showing higher levels of EGR1 expression in the IL (U = 8.00, P< 0.05). Finally, we compared the ratio of PL to IL activity between males and females using a Mann-Whitney U test (Figure 4D). There was a significant difference between groups (U = 0.00, P < 0.001) with males showing higher ratios than females.

**Figure 4.**
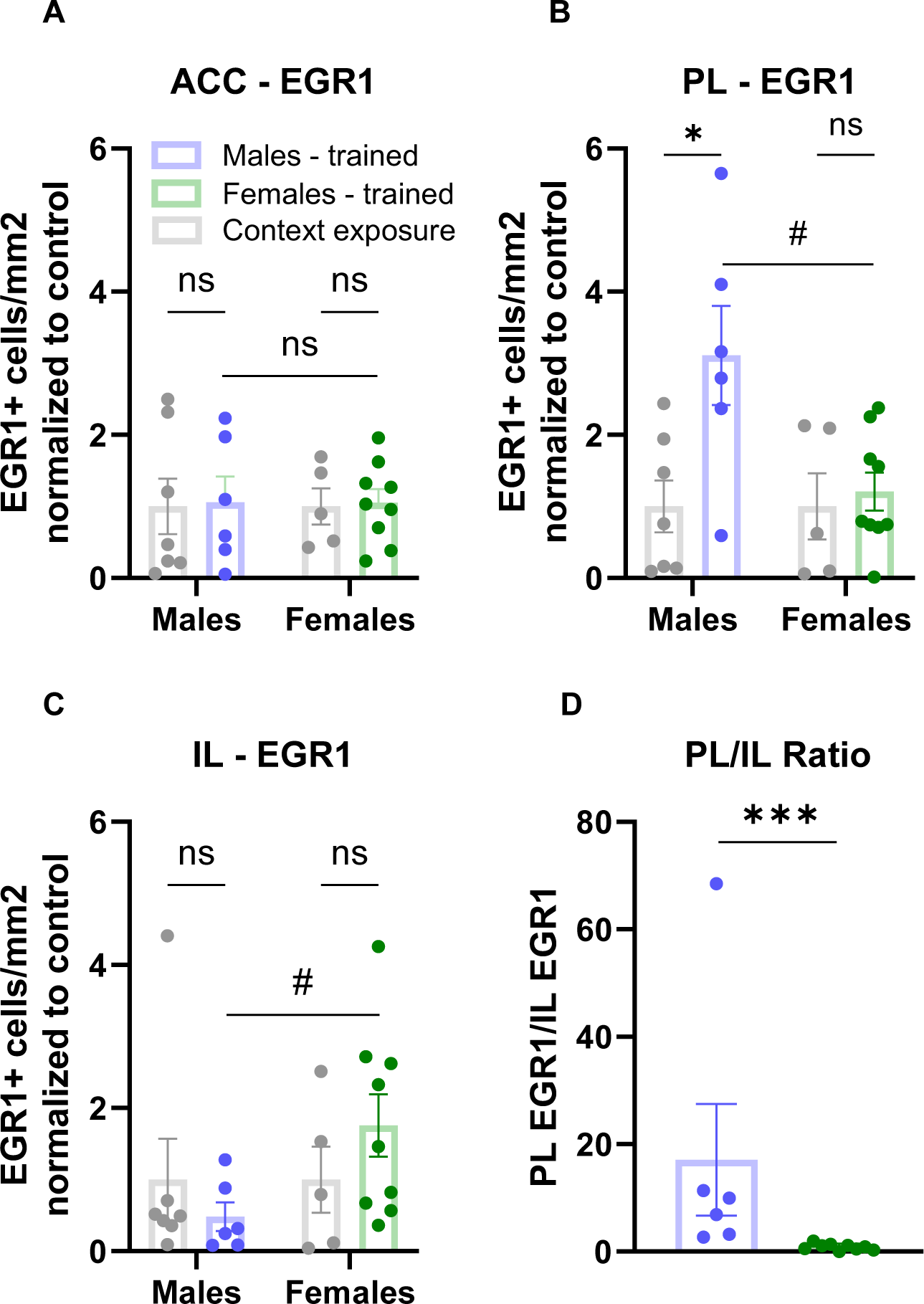
The number of EGR1-positive cells in neurons not expressing AAVretro in the ACC (A), PL (B) , and IL (C) following the expression of contextual fear. (D) The average ratio of the EGR1-positive cells in the PL to ERG1-positive cells in the IL for males and females that received conditioning and testing. * p < 0.05, *** p < 0.001.

## Discussion

This study sought to determine if projections from the medial prefrontal cortex to the ventrolateral periaqueductal gray were activated by the expression of contextual fear and whether sex differences in contextual fear were associated with neural activity in mPFC-vlPAG projections. We also assessed the pattern of neural activity in the mPFC in cells which were not labeled by the retrograde tracer. Male rats exhibited higher levels of freezing than females during the test session, consistent with several studies [3, 4, 7, 10]. Tracer injections in the vlPAG produced dense labeling ACC, PL, and IL. The general pattern of retrograde expression in the mPFC was similar in males and females and was generally consistent with prior tracing studies [34]. The level of EGR1 expression in prelimbic projections to the vlPAG was significantly higher than controls in male rats, but there was no significant difference between trained females and control females. However, there was no significant difference when the sexes were compared directly, indicating the lack of a sex difference. There were no differences in the expression of EGR1 between groups in males or females in either the ACC or IL projections to the vlPAG. When we counted EGR1 positive cells in the same three regions of the mPFC in neurons not labeled by the tracer we found increased expression in the PL in males, but not females, and significantly higher expression in males when the sexes were compared directly. In the IL cortex, neither trained male nor female rats were different than controls, however trained females showed higher levels of EGR1 expression than males. There were no differences in EGR1 expression in the ACC between groups or between sexes.

Our findings in male rats are consistent with some prior studies outlining a role for the prelimbic cortex in the expression of fear [13, 26, 35 36]. Nearly all the prior work characterizing the role of the PL in fear expression has been completed in males, and while some studies have demonstrated sex differences in mPFC function related to various aspects of fear processing [18–21] to our knowledge no prior studies have compared neural activity markers in males and females associated with the expression of contextual fear. Our results suggest that there are notable differences between males and females. First, while males show increased activation of the PL following contextual fear expression females do not. Second, while there was no change in activation of the IL following the expression of contextual fear in males, female rats show increased activation of IL relative to males. We also examined the relative activity levels of the PL and IL, as the prior work suggested that the balance of activity between the PL and IL is predictive of fear expression [17]. We found that males exhibited a much higher level of PL activity relative to IL than females. Together, these findings indicate that sex differences in the expression of contextual fear may involve differences in the activity levels of the PL and IL.

The increased activation of the IL in females might suggest, counterintuitively, that the IL drives fear expression in females. However, given the prior work tying IL to fear inhibition, it is perhaps more likely that the increased expression of EGR1 in IL is antecedent to reduced expression of contextual fear in females. While the role of IL in fear inhibition has been studied primarily in the context of extinction learning and the recall of extinction, recent work provides an indication that IL may participate more broadly in the inhibition of fear. Studies in male rats have reported that there is a suppression of neural activity in the IL after contextual fear conditioning [37] and cued fear conditioning [17], and that the suppression of IL activity may underlie the expression of fear [17]. Consistent with this, earlier work showed that pharmacological activation of IL prior to testing disrupted the expression of contextual fear and facilitated the subsequent extinction [16]. Similarly, post-training activation of IL was recently shown to disrupt the expression of cued fear and enhance the subsequent extinction [38]. In the latter study, similar effects were obtained when IL activation occurred 24 hours after training, but 24 hours before testing, indicating that the effects of IL stimulation are not necessarily the result of disrupted memory consolidation or an acute effect on the expression and extinction of cued fear, but rather may be enhancing an inhibitory memory. Considering these recent findings, one plausible explanation of our results is that the reduced expression of contextual fear in females and increased activation of the IL reflects the expression of an inhibitory memory. If this is the case, it will be important to determine why we did not observe increased IL activity in males, as some of these recent studies showing long-term effects of IL stimulation were performed in both sexes [38]. One possibility is that when conditioning and the expression of fear occur without manipulation of neural activity, females form a stronger IL-based inhibitory memory than males and that activating the IL in males essentially mitigates this sex difference.

In male rats we found that projections from the PL to the vlPAG were activated following the expression of contextual fear. Levels of EGR1 in PL projections to the vlPAG in females fear conditioned and tested were not different than controls, however a direct comparison of trained males and females was not significant indicating the absence of a robust sex difference. Overall levels of activity in the PL did differ between sexes, so if the sex differences in contextual fear expression does not involve PL inputs to vlPAG then some other pathway must be involved. At least two pathways are of interest, including PL projections to the basolateral amygdala and to the paraventricular thalamus, both of which have been implicated in fear expression in males [26]. Our finding in males that PL-vlPAG is active following fear expression is in apparent contrast with a prior study [31], which revealed that dorsomedial prefrontal (i.e. ACC and PL combined) projections to the vlPAG are not involved in the expression of contextual fear but rather the discrimination of contextual fear. This study showed that inactivation of dmPFC-vlPAG projections prevented the reduction of fear when rats were transferred to a safe context, whereas activation of this pathway reduced freezing behavior when rats were tested in the training context. One key difference that might explain the discrepancies is that we were able to separate PL and ACC, whereas the prior work did not distinguish PL from ACC, or lateral PAG from ventrolateral PAG. Given the large number of studies describing functional differences within discrete areas of the mPFC [39, 40] and PAG [41, 42], it is likely that there is specialization of function in mPFC projections to the PAG.

Recent studies have begun to address the question as to whether and how the neural mechanisms underlying fear expression in females differ from those in males. A study in mice [11] found that the expression of contextual fear was associated with sex differences in cFos expression in the dorsal hippocampus and amygdala. In females, the expression of fear was associated with increased neural activity in the CA1 region of the hippocampus, while males showed increased activation in CA3 and the dentate gyrus, in addition to CA1. In the amygdala, females showed higher levels cFos in the basal amygdala whereas no change was detected in males. Another recent study [43] examined cFos activity in the amygdala and bed nucleus of the stria terminalis (BNST) following contextual fear expression in male and female rats. This study reported that males, but not females, showed higher levels of neural activity in the BNST. Both sexes showed activation in the lateral amygdala following context fear expression, however somewhat surprisingly there was no change in cFos in either sex in either the central or basal amygdala. These published studies and our findings reported here point to substantive sex differences in the neural circuits mediating contextual fear expression. However, integrating these studies is complicated by the fact that the behavioral results are not consistent across studies. The former study reported higher levels of contextual fear expression in females, whereas the latter found no sex difference, although a trend for higher freezing in males was noted. Prior studies have identified a variety of factors mediating sex differences in contextual fear [8–10, 46] but more comprehensive behavioral work is necessary to identify additional factors and how they influence the neural mechanisms supporting contextual fear.

In summary, the results from these experiments demonstrate noteworthy sex differences in neural activity in the prefrontal cortex associated with the expression of contextual fear. In prefrontal projections to the ventrolateral PAG, the expression of contextual fear was associated with the activation of PL (but not ACC or IL) projections in male rats, whereas in females there was no change in activity of the projections from any region of the mPFC to the vlPAG. Outside of the projections to vlPAG, we observed that males exhibited higher levels of neural activity in the PL following fear expression than did females. In the IL, females showed higher levels of activity associated with fear expression than did males, suggesting that the reduction in the expression of contextual fear in females may be the result of increased inhibition by IL of the neural circuits that drive fear expression.

## Author Contributions

R.G.P and K.V. both collected and analyzed the data and drafted the manuscript.

